# A miR-494 dependent feedback loop regulates ER stress

**DOI:** 10.1101/2020.05.12.088856

**Authors:** Namita Chatterjee, Cristina Espinosa-Diez, Sudarshan Anand

## Abstract

Defects in stress responses are important contributors in many chronic conditions including cancer, cardiovascular disease, diabetes, and obesity-driven pathologies like non-alcoholic steatohepatitis (NASH). Specifically, endoplasmic reticulum (ER) stress is linked with these pathologies and control of ER stress can ameliorate tissue damage. MicroRNAs have a critical role in regulating diverse stress responses including ER stress. Here we show that miR-494 plays a functional role during ER stress. ER stress inducers (tunicamycin & thapsigargin) robustly increase the expression of miR-494 *in vitro* in an ATF6 dependent manner. Surprisingly, miR-494 pretreatment dampens the induction and magnitude of ER stress in response to tunicamycin in endothelial cells. Conversely, inhibition of miR-494 increases ER stress *de novo* and amplifies the effects of ER stress inducers. Using Mass Spectrometry (TMT-MS) we identified 23 proteins that are downregulated by both tunicamycin and miR-494. Among these, we found 6 transcripts which harbor a putative miR-494 binding site. We validated the anti-apoptotic gene *BIRC5* (survivin) as one of the targets of miR-494 during ER stress. Finally, induction of ER stress *in vivo* increases miR-494 expression in the liver. Pretreatment of mice with a miR-494 plasmid via hydrodynamic injection decreased ER stress in response to tunicamycin in part by decreasing inflammatory chemokines and cytokines. In summary, our data indicates that ER stress driven miR-494 may act in a feedback inhibitory loop to dampen downstream ER stress signaling. We propose that RNA-based approaches targeting miR-494 or its targets may be attractive candidates for inhibiting ER stress dependent pathologies in human disease.

## Introduction

The endoplasmic reticulum (ER) is the site of mRNA translation alongside proper folding and post-translational modifications of proteins destined for secretion or localization to various cellular membrane systems. In addition, the ER is the center of lipid biosynthesis, detoxification, homeostasis of intracellular Ca^2+^ and redox balance. Several pathologies including neurodegenerative diseases, diabetes, atherosclerosis and cancers have attributed ER stress as a critical driver of disease. When the folding capacity of the ER is challenged, the accumulation of un- or mis-folded proteins in the lumen of the ER triggers the unfolded protein response (UPR) (1,2). This response activates a trio of transducers: the PKR-like ER kinase (PERK), inositol requiring enzyme 1 α (IRE1α) and activating transcription factor 6 (ATF6), which work synergistically to control transcriptional and translational programs to either alleviate the burden of unfolded proteins and return to protein homeostasis or initiate apoptosis. It is thought that acute ER stress can trigger feedback mechanisms that protect cells from death by suppressing global translation and increasing ER chaperone levels, whereas persistent UPR activation or chronic unmitigated ER stress leads to increased oxidative stress, inflammation and eventual apoptosis (3,4).

Endothelial cells (ECs) encounter a variety of stressors and stimuli during development and disease (5). Recent studies have implicated ER stress and the UPR as drivers of endothelial dysfunction in cardiovascular disease (6,7). For instance, ER stress is thought to promote both EC inflammation and apoptosis in atherosclerosis (8–10). Furthermore, ER stress has been shown to contribute to vascular dysfunction and cardiac damage in preclinical models of hypertension (11). Similarly, oxidative stress and ER stress pathways have been shown to be interlinked in ECs in metabolic syndromes such as diabetes and non-alcoholic fatty liver disease (NAFLD) (12–16). Therefore, ECs have adopted several mechanisms to regulate cell fate decisions in response to both acute and chronic ER stress (17).

MicroRNAs (miRs) are small non-coding RNAs that are critical regulators of physiological and pathological stress responses (18,19). We, and others have shown that specific miRs regulate EC responses to numerous effectors including: angiogenic growth factors (20,21), hypoxia (22,23), DNA damage (24), and oxidative stress (25). Furthermore, specific miRs have been identified as modulators of ER stress in different models of disease (26–29). Indeed Kassan *et al*., identified miR-204 as a promoter of ER stress in ECs by targeting the SIRT1 pathway (30). Our previous work identified a cohort of miRs that are induced by DNA damage in ECs (24,31,32). We further characterized one of these miRs, miR-494, as a regulator of endothelial senescence in response to genotoxic stressors (24). In this study, we show that ER stress is a potent inducer of miR-494 likely via ATF6. Surprisingly, we find that miR-494 operates in a feedback loop to dampen the ER stress response potentially by targeting the anti-apoptotic protein survivin. Our *in vivo* experiments suggest miR-494 pretreatment diminishes tunicamycin-induced ER stress in the liver. Overall, our studies illuminate a novel function for miR-494 and open new avenues for further investigations into mechanisms by which miRs modulate stress responses.

## Methods

### Cell culture and reagents

HUVECs (Cat: NC9946677, Lonza) were cultured in EGM-2 media (Cat: CC-3162, Lonza) supplemented with 10% fetal bovine serum (Cat: FB-21, Lot# 649116, Hyclone) and were maintained at 37°C with 5% CO_2_. Cells used for experiments were low passage number (3–8). Tunicamycin (Cat: T7765, Sigma) was dissolved in DMSO (Sigma).

### Cell transfection

HUVECs (50% confluence) were transfected with 50nM mimic/inhibitor RNA using the lipofectamine RNAiMax reagent (Cat: 13778-150, Invitrogen) according to the manufacturer’s instructions. Specifically, mirVana miR mimic negative control (Cat: 4464061, Ambion), hsa-miR-494-3p mimic (MIMAT002816) (Cat 4404066, Assay ID MC12409, Ambion), mirVana miR inhibitor negative control #1 (Cat 4464076, Ambion), hsa-miR-494-3p inhibitor (MIMAT002816) (Cat 4464084, Assay ID MH12409, Ambion). Negative Control LNA mimic (Cat: YM00479903, Qiagen), hsa-miR-494-3p miRCURY LNA™ microRNA Mimic (YM00472106, MIMAT002816, Qiagen), miR-control (CAT#4464058), miR-494 inhibitor (CAT#4464084/MIMAT0004975), and miR inhibitor negative control (CAT#4464076) were purchased from Ambion/Life Technologies.

For gene knock down, HUVECs were transfected with 10nM siRNA using the lipofectamine RNAiMax reagent (Cat: 13778-150, Invitrogen) according to the manufacturer’s instructions. Specifically, siRNA for ATF6 (Cat: SR307883A, Origene), EIF2AK3 (Cat: SR306267A, Origene), ERN1 (Cat: SR301457A&B, Origene), siRNA Negative Control (Cat: SR30001, Origene).

### Gene expression

Total mRNA was isolated from cells and tissues using the miRNeasy Mini Kit (cat: 217004, Qiagen). Reverse transcription was performed using High Capacity cDNA Reverse Transcription Kit (Cat: 4368814, Applied Biosystems) according to the manufacturer’s instructions. Gene expression was measured using real-time quantitative PCR (qRT-PCR) with TaqMan™ Master Mix II no UNG (Cat 4440049, Thermofisher Scientific) with the following primer/probe sets: miR-494-3p (Cat: 4427975, Assay ID: 002365), human primary miR-494 (Cat: 4427013, Assay ID: Hs04225959_pri) U6 snRNA (Cat: 4440887, Assay ID: 001973), sno234 (Cat: 4440887, Assay ID: 001234) GAPDH (Cat 4351370 Assay ID: Hs02758991_g1) according to the manufacturer’s instructions. SYBR Green qRT-PCR assays were conducted using PowerUp SYBR Green Master Mix (Cat A25741, Thermofisher Scientific) with primers shown in Table 1. Fold change was calculated using the 2^−ΔΔCt^ method relative to an internal control (GAPDH, β-Actin (ActB), sno234 or U6).

**Table 1:**
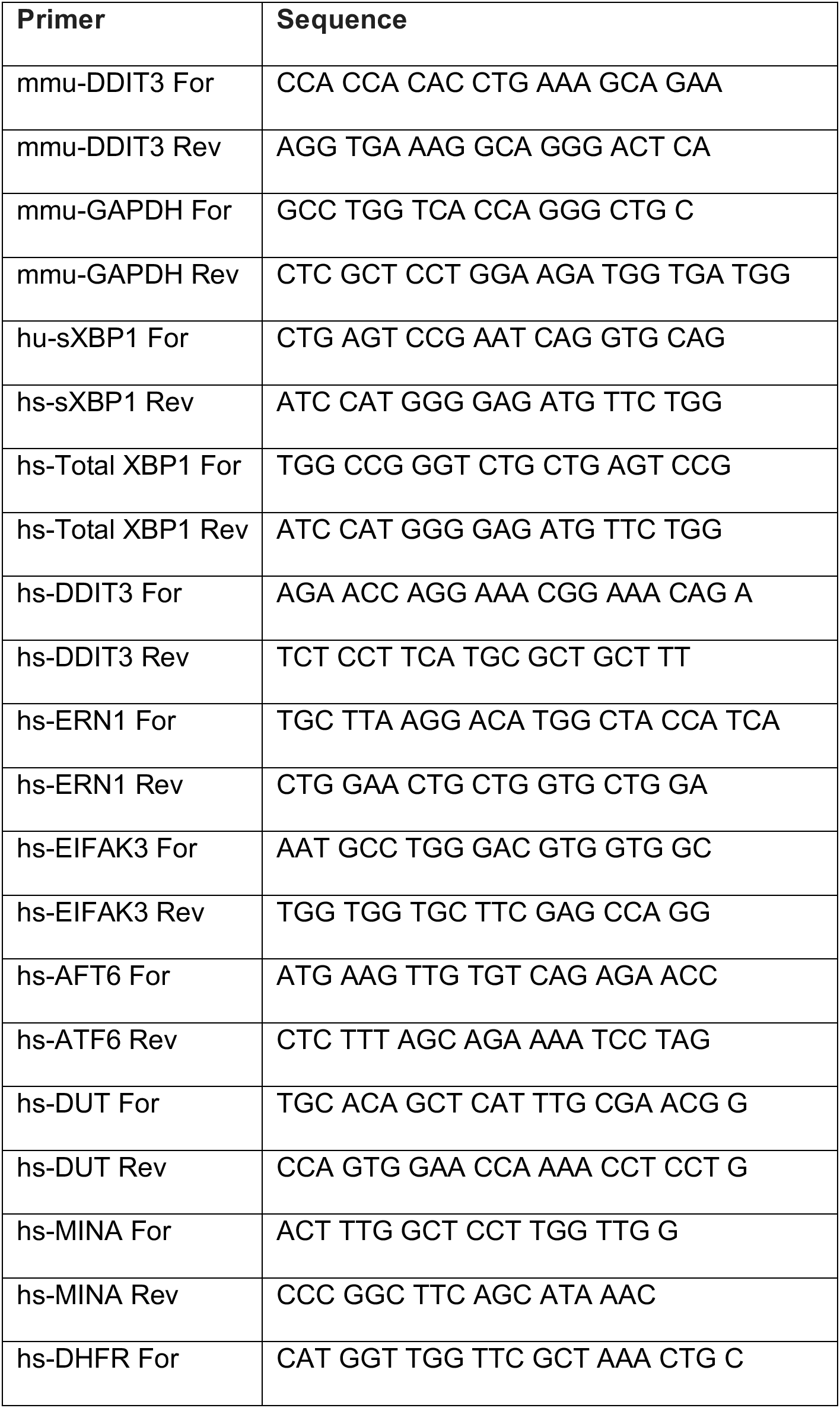
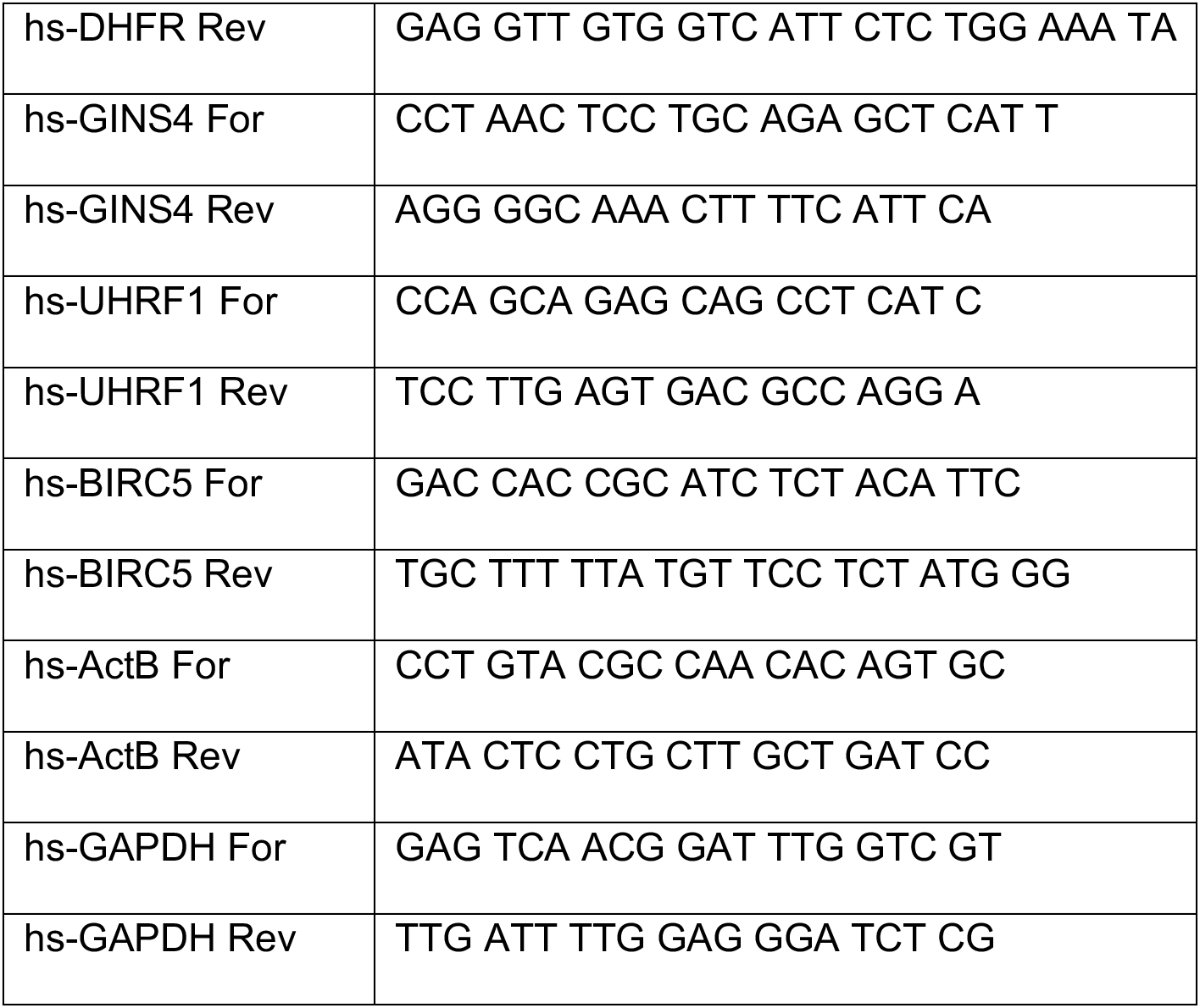
Oligonucleotide primers used for SYBR Green based qRT-PCR assays

### Nanostring^®^ Immune Profiling

We used the Nanostring PanCancer Mouse IO360™ Gene Expression Panel containing 770 genes involved in the tumor, surrounding microenvironment and immune response in cancer. Livers were homogenized and total RNA was isolated using the miRNeasy Mini Kit (cat: 217004, Qiagen). 100ng of RNA was used to perform the generate the expression profile as per the manufacturer’s instructions. Data was analyzed using the nSolver™ Analysis software. Raw data was normalized to a panel of housekeeping genes.

### Immunofluorescence

For immunofluorescence analysis, HUVECs were seeded (12,000 cells/well) on coverslips in a 24-well plate and transfected with mimic/inhibitor for 24h as described above. The following day, the media was aspirated and replaced with fresh media containing tunicamycin (10μg/mL TCN, Sigma) or DMSO (Sigma) for 24-48h. The media was aspirated and cells were rinsed in ice cold PBS for 2 min followed by fixation in 4% paraformaldehyde diluted in PBS (Cat: NC9658705, FisherSci) for 10 min at room temperature (RT). Coverslips were rinsed in PBS three times and incubated in serum free DAKO Protein Block (X0909, DAKO) for 1h at RT. Next, coverslips were incubated overnight at 4°C in anti-survivin primary antibody diluted in PBS/5% BSA. After rinsing in PBS three times, the coverslips were incubated in the dark with Alexa Fluor^®^ 488 goat anti-rabbit secondary antibody diluted in PBS/5% BSA. Refer to table 2 for antibody information and dilutions. After rinsing in PBS three times, the coverslips were mounted onto glass slides with Aqua-Poly/Mount (Cat No. 18606, Polysciences Inc). Slides were left to cure overnight at RT prior to imaging. Coverslips were imaged using a Nikon/Yokogawa CSU-W1 Spinning Disk Confocal Microscope. All imaging settings remained constant throughout the imaging sessions. The percentage stained area was analyzed using the open-access Image J software (NIH).

**Table 2:** Antibodies used in this study

| Antibody | Source | Identifier | Concentration |
| --- | --- | --- | --- |
| Survivin | Cell Signaling Technology | 2808 | 1:25 |
| CHOP | Novus | NB600-1335 | 1:250 |
| XBP1-s | Cell Signaling Technology | 83418 | 1:25 |
| GINS4 | Novus | NBP2-16659 | 1:25 |
| Alexa Fluor® 488<br>(goat anti-rabbit) | Cell Signaling Technology | 4412 | 1:100 |

### Simple western blot

HUVECs were seeded in 6 well plates (200,000 cells/well) and transfected as described above. After 24h, cells were treated with 10μg/mL TCN for 48h. After treatment, the media was aspirated and the cells were washed twice in ice cold PBS and lysed directly in the plate in RIPA buffer (cat: PI89900, Pierce) containing Protease Inhibitor Mini Tablets (1 tablet/10mL RIPA buffer, Cat: A322953, Pierce) with phosphatase inhibitor cocktail 2 & 3 (1:1000, Cat: P5726 and P0044) for 15 min on ice. Lysates were rotated at 4°C for 30 min-2h and then centrifuged at 12,000 × g at 4°C for 40 min. The supernatant was collected and protein concentration was determined using the Pierce BCA Protein assay kit (Cat No. 23225). Samples were diluted to 0.75μg/mL with 1× Sample Buffer (ProteinSimple). Protein quantification was performed using a 12–230 kDa 25 lane plate (PS-MK15; ProteinSimple) in a ProteinSimple Wes Capillary Western Blot analyzer according to the manufacturer’s instructions. The standard Simple Western protocol was altered to increase sample load time to 13s and a separation time of 33 min.

### Animal experiments

For *in vivo* experiments, male C57Bl/6 mice (aged 12-20 weeks) were purchased from Jackson labs. Animals were housed in a controlled animal facility with access to normal rodent chow and water *ad libitum*. All experimental protocols were approved by the Institutional Animal Care and Use Committee (IACUC) at Oregon Health and Science University (OHSU). For *in vivo* studies, the LentimiRa-GFP-hsa-miR-494-3p Vector (Cat: mh10739, Applied Biological Materials, abmgood.com) and pLenti-III-miR-GFP Control Vector (Cat: M001, Applied Biological Materials) were purified using the EndoFree Plasmid Maxi Prep Kit (Cat: 118379, Qiagen). Plasmids (20-30μg in PBS) were injected using a hydrodynamic injection method into the tail vein with a volume of 1.5-2mL/animal in 7-10s as described (33,34). After 24h, animals were injected *i.p.* with tunicamycin (2 μg/g Cat: T7765, Sigma) which was prepared by dissolving the compound in a small volume of DMSO (100μL, Sigma) and then diluting in 150mM Sucrose (Cat: 57-50-1, FisherSci) in dH_2_O. After 6-20h, livers were harvested and total RNA was isolated using the miRNeasy Mini Kit (Cat: 217004, Qiagen).

### TMT-Mass Spectrometry

TMT labeling and mass spectrometry were performed by the OHSU proteomics core facility as described in detail elsewhere (35). Briefly, HUVECs were treated with microRNAs or tunicamycin as described above. After 24h-48h, samples were lysed in 50 mM triethyl ammonium bicarbonate (TEAB) buffer (50 μg of protein/ sample) followed by a Protease Max digestion, a microspin solid phase extraction and TMT labeling. Multiplexed TMT-labeled samples were separated by two-dimensional reverse-phase liquid chromatography. Tandem mass spectrometry data was collected using an Orbitrap Fusion Tribrid instrument (Thermo Scientific). RAW instrument files were processed using Proteome Discoverer (PD) (Thermo Scientific). Searches used a reversed sequence decoy strategy to control peptide false discovery and identifications were validated by Percolator software. Search results and TMT reporter ion intensities were processed with in-house scripts. Differential protein abundance was determined using the Bioconductor package edgeR.

### Statistical Analysis

All statistical analysis was performed using Prism software (GraphPad Software, San Diego, CA). Differences between pairs of groups were analyzed by Student’s *t*-test. Comparison among multiple groups was performed by one-way ANOVA followed by a post hoc test (Tukey’s or Holm-Sidak). In the absence of multiple comparisons, Fisher’s LSD test was used. Values of *n* refer to the number of experiments used to obtain each value. For mouse studies where the data was not normally distributed, we used two-tailed Mann-Whitney U test. Values of *p* ≤ 0.05 were considered significant

## Acknowledgements

This work was supported by funding from NHLBI to S.A. (R01 HL137779 and R01 HL143803). We thank Drs Larry David, Ashok Reddy, and John Klimek of the OHSU Proteomics core facility for Mass Spectrometry experiments. Mass spectrometric analysis was performed by the OHSU Proteomics Shared Resource with partial support from NIH core grants P30EY010572 P30CA069533 and S10OD012246. We thank Ms. Rebecca Ruhl for technical help and members of the Anand lab for useful discussions.

## Author Contributions

N.C, C. E-D and S.A. designed experiments, N.C, C.E-D performed experiments, analyzed the data, N.C and S.A. wrote the manuscript.

## Results

### ER stress induces miR-494 *in vitro*

We previously showed that miR-494 is responsive to radiation and chemical inducers of genotoxic stress and functions to increase endothelial senescence during DNA damage responses (24). Given the intricate relationship between radiation, oxidative stress and ER stress (14,17), we asked if ER stress affected miR-494 expression and function. First, we confirmed a robust ER stress response to known inducers, tunicamycin (TCN) and thapsigargin (TG), in human umbilical vein endothelial cells (HUVECs) by measuring the level of the transcription factors spliced *XBP1* (mRNA: *sXBP1*, protein: XBP1s) and *DDIT3* (CHOP), which are well characterized markers of the ER stress response (Fig. 1A-B). We observed that TCN significantly increased the levels of mature miR-494 (Fig. 1C) and to a lesser extent, the primary unprocessed miR-494 transcript (Fig. 1D). Similarly, TG also induced *sXBP1* and *DDIT3* in parallel with the primary and mature forms of miR-494 (Supplementary Fig. 1). We also saw a comparable increase in both ER stress and the primary miR-494 transcript in another EC line (Human Microvascular Endothelial cells - HMVECs) (Supplementary Fig. 2).

**Figure 1:**
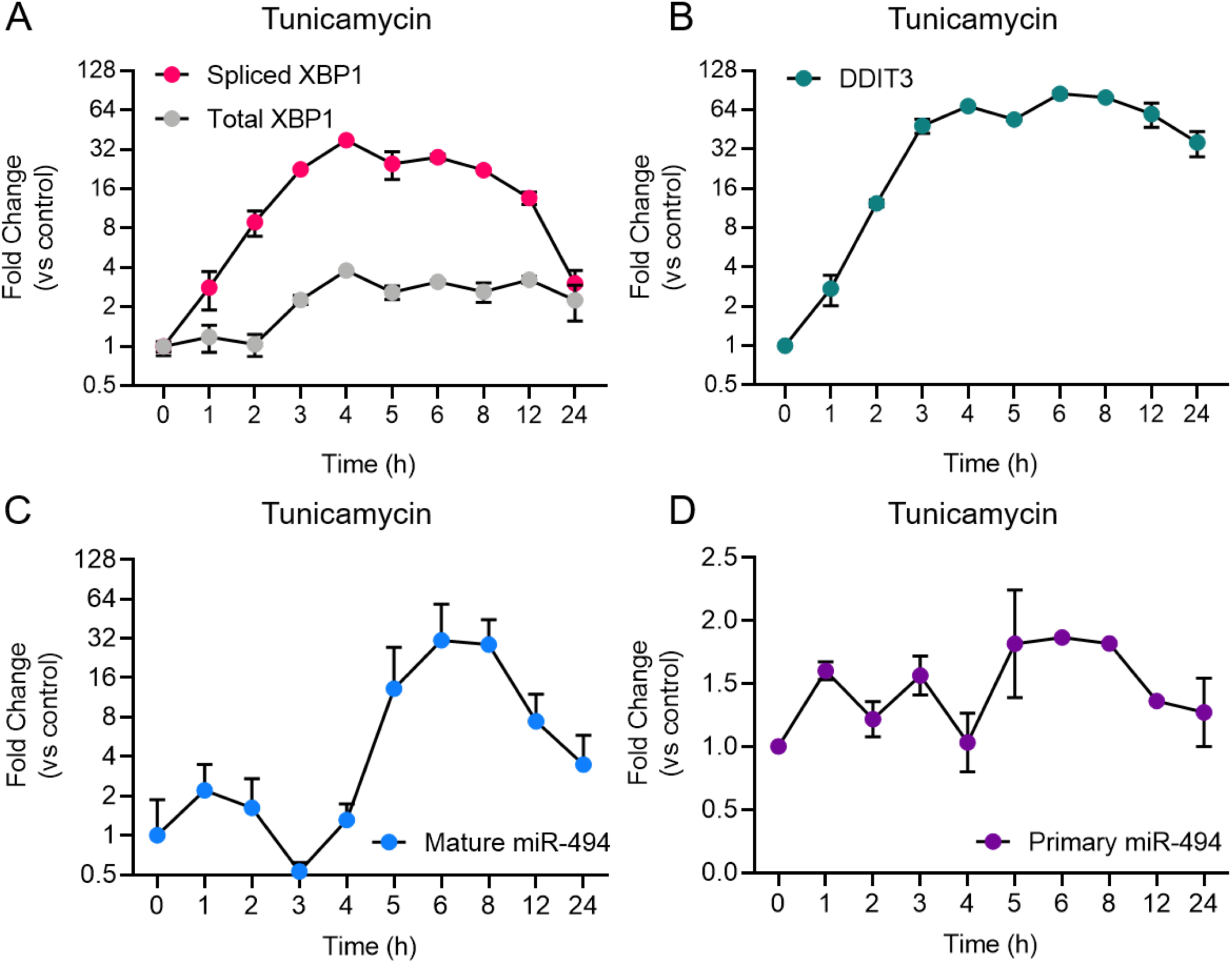
ER stress induces expression of the primary and mature forms of miR-494. Relative mRNA expression of ER stress responsive genes as measured using qRT-PCR. A) spliced XBP1 and Total XBP1 and B) *DDIT3* (CHOP) levels in HUVECs treated with 5μg/mL tunicamycin (TCN) over a time course. Relative expression of C) primary miR-494 (pri-miR-494) and D) mature miR-494 in HUVECs treated with TCN over a time course. Gene expression is normalized to GAPDH or U6 and mean fold change compared to vehicle control or time 0h is shown. Graphs are representative of one biological replicate from three independent replicates where values indicate mean ± standard deviation.

A triumvirate of transducers (ATF6, IRE1α & PERK) co-ordinate ER stress response programs to promote cell recovery, or in the case of excessive or chronic stress, initiate apoptosis. We asked which of these transducers was responsible for miR-494 induction in response to ER stress using siRNAs. We found that knockdown of ATF6 but not IRE1α (*ERN1*) or PERK (*EIF2AK3*) significantly decreased the induction of miR-494 6h after TCN treatment (Supplementary Fig. 3). These data indicate that ER stress induces the expression of miR-494 in an ATF6 dependent manner.

### miR-494 diminishes ER stress

To understand the functional relevance of miR-494 during ER stress, we performed gain- and loss-of function experiments utilizing miR-494 mimics or inhibitors, respectively. Pretreatment of HUVECs with miR-494 mimic suppressed the TCN based induction of ER stress responsive genes *DDIT3* (Fig. 2A). Conversely, inhibition of miR-494 robustly increased levels of *DDIT3* (Fig. 2B). Similarly, miR-494 mimic decreased levels of *sXBP1* (Fig. 2C) whereas, inhibition of miR-494 increased sXBP1 mRNA both *de novo* and in combination with TCN (Fig. 2D). We validated that the miR-494 mediated decrease in these transcription factors also persisted at the protein level (Fig. 2E). Taken together, our data suggest that miR-494 diminishes the induction of ER stress-associated transcription factors and plays a protective role in HUVECs during ER stress.

**Figure 2:**
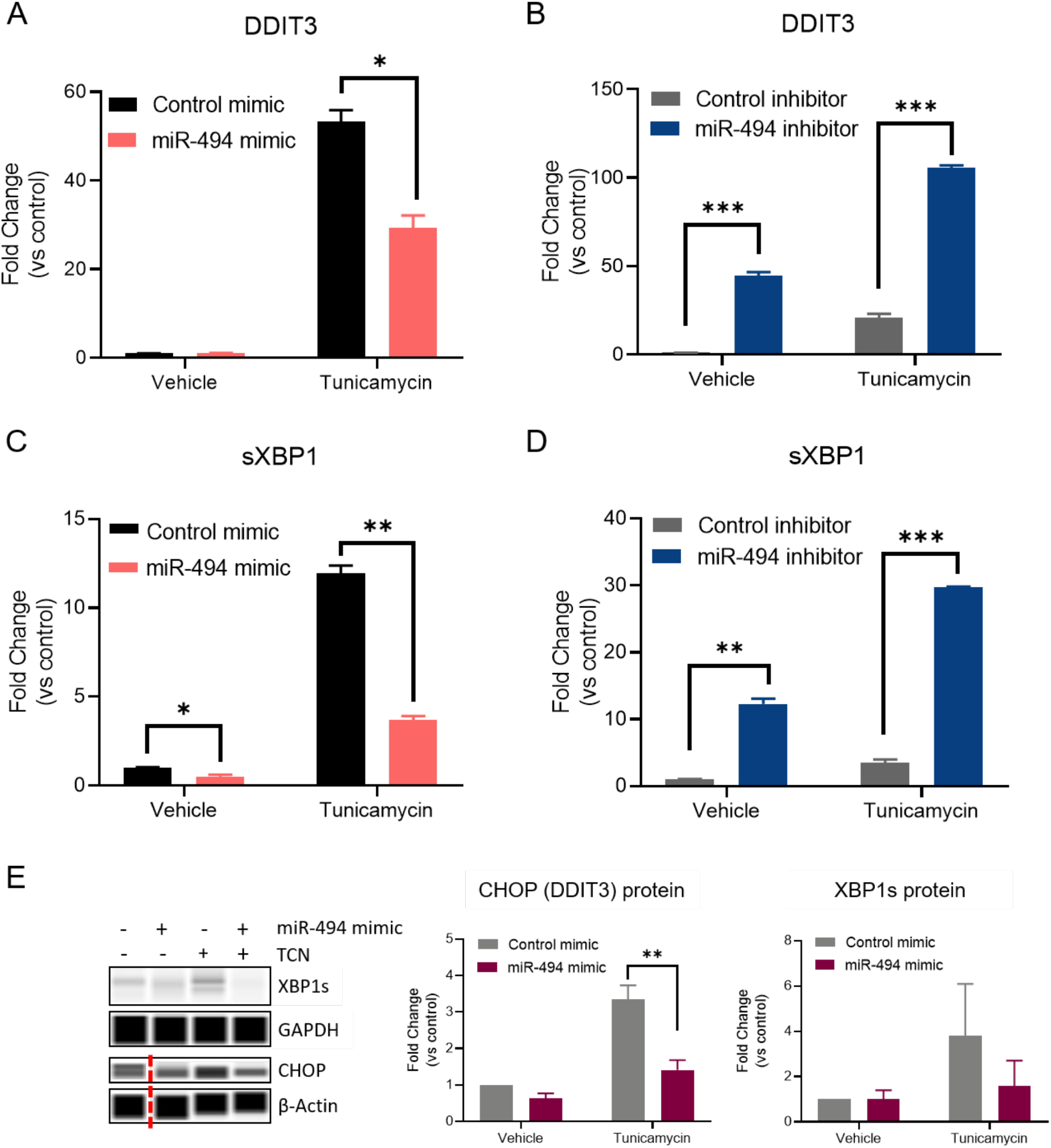
miR-494 is a negative regulator of ER stress *in vitro*. Relative mRNA expression of ER stress responsive genes as measured by qRT-PCR. A-B) *DDIT3* (CHOP), C-D) spliced XBP1 in HUVECs treated with 10μg/mL TCN 48h after transfection with A, C) miR-494 mimic or B, D) miR-494 inhibitor. Gene expression is normalized to GAPDH and mean fold changes compared to control treatments are shown. E) Simple Western blot analysis of HUVECs transfected with miR-494 mimic or control (24h) followed by TCN (10μg/mL) for 24h. Vertical dotted red line indicates non-adjacent lanes. Graphs are mean + SEM fold changes of biological replicates from *n*=3 independent experiments. * P<0.05, ** P< 0.01, ***P<0.001 by two-tailed Student’s T-test.

**Figure 3.**
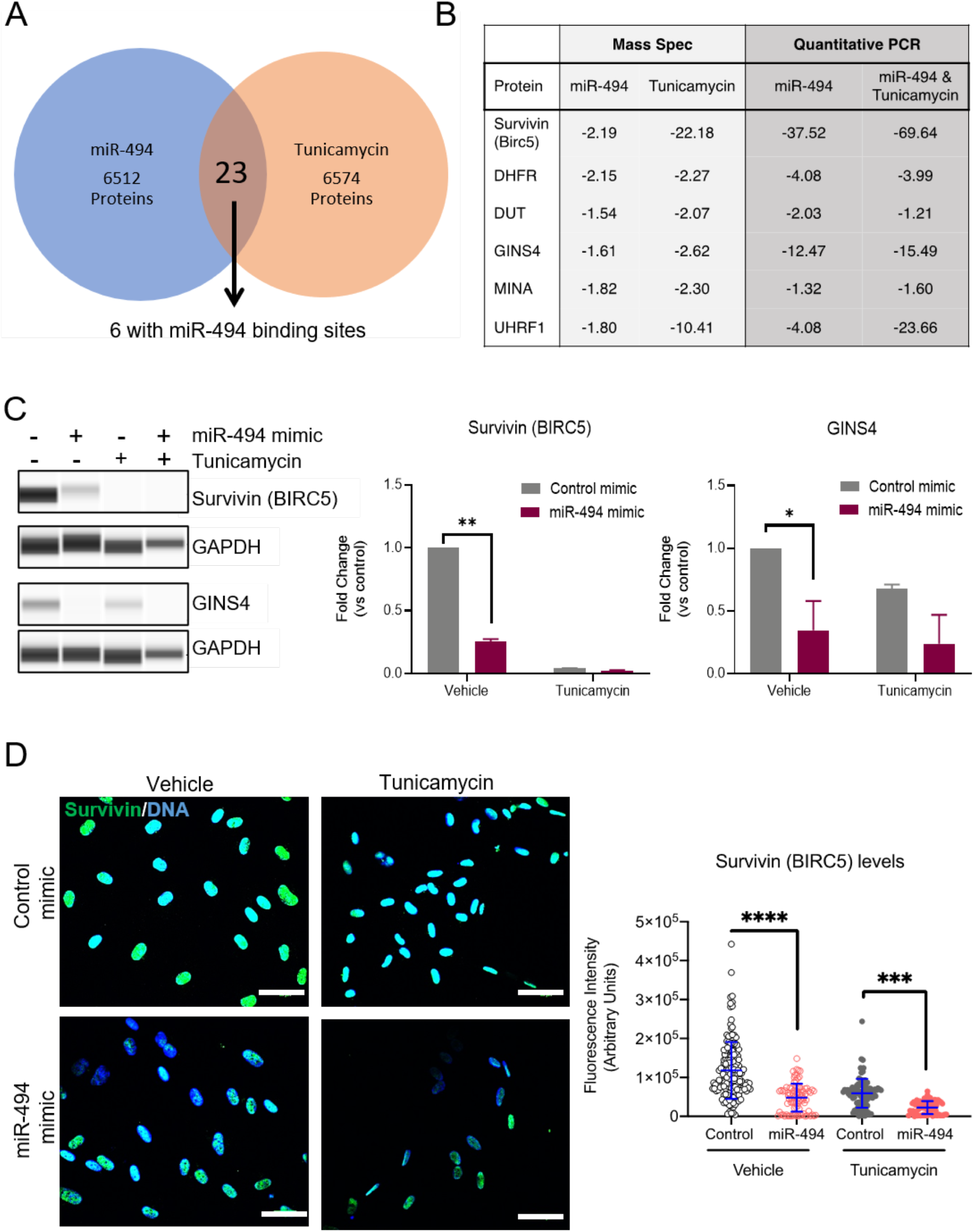
miR-494 regulates target genes in cell survival and DNA replication. A) Venn diagram showing the number of downregulated target proteins in a Tandem Mass Tag labelled Mass Spectrometry profile from HUVECs treated with TCN or transfected with miR-494 compared to vehicle treatment or control miR respectively. B) Fold-change (compared to respective controls) of protein or mRNA levels as assessed by Mass Spectrometry or qRT-PCR respectively for the 6 targets that were downregulated in both TCN and miR-494 groups. All 6 targets harbor miR-494 binding sites in their 3’UTRs. C) Representative Simple Western blot showing survivin and GINS4 levels in HUVECs 24h after miR-494 transfection followed by TCN treatment (24h). Right panels show quantitation of biological replicates. * *p*<0.05, ** *p*< 0.01, by two-tailed Student’s T-test. D) Immunofluorescence imaging showing survivin expression in HUVECs 24h after miR transfection and/or TCN treatment. Right panel shows Survivin fluorescence intensity quantifications via Image J in each cell. Each dot represents individual cells. Scale bar in white = 50 μm. * P<0.05, ** P< 0.01, ***P<0.001 by one-way ANOVA with post-hoc Tukey’s correction.

### Mass spectrometry identifies putative targets of miR-494 relevant for ER stress

miRs typically are thought to regulate numerous targets in a context dependent manner. We have previously shown that miR-494 targets the Mre11a, Rad50, Nbn (MRN) complex in the DNA damage repair pathway in response to genotoxic stressors (24). To identify the targets relevant for miR-494 in the context of ER stress, we undertook a proteomics-based approach. We used Tandem mass tag-based mass spectrometry (TMT-MS) to compare changes in the proteome with either miR-494 mimic treatment or TCN treatment. We found 23 proteins were commonly downregulated between the TCN and miR-494 treatment conditions (Fig. 3A).

Among these, we identified six proteins whose mRNAs harbored putative miR-494 binding sites in their 3’UTRs as predicted by the TargetScan algorithm. We validated the expression of these six genes using qRT-PCR (Fig. 3B and Supplementary Fig. 4A-D). All six targets were validated and demonstrated a substantial decrease in expression in response to miR-494 gain-of-function activity. Conversely, inhibition of miR-494 restored *BIRC5*, *GINS4, MINA* and to a lesser extent, *UHRF1* mRNA levels in HUVECs treated with TCN (Supplementary Fig 4E). Since *BIRC5* and *GINS4* were consistently regulated in a miR-494 dependent fashion, we chose to further validate these two targets at the level of protein expression. We found survivin (gene: *BIRC5*), and GINS4 protein levels were significantly decreased upon treatment with miR-494 mimic alone, TCN treatment alone, and the sequential combination of miR pre-treatment (24h) followed by TCN (24h) (Fig. 3C). Finally, we used immunofluorescence staining and confocal microscopy to evaluate survivin expression in HUVECs. Consistent with our western blot data, we observed a significant decrease in the amount of survivin in cells transfected with either miR-494 mimic or TCN treatment alone or in the sequential combination (Fig. 3D). Multiple studies have demonstrated that miR-494 binds to the 3’UTR and leads to subsequent degradation of the survivin (*BIRC5*) transcript (36–38). Our observations support these data and further demonstrates this relationship in ECs during ER stress.

### miR-494 pretreatment ameliorates ER stress in mouse liver

ER stress is thought to be a significant driver of liver diseases including hepatic fibrosis, non-alcoholic fatty liver disease (NAFLD) that can progress to steatohepatitis (NASH) and hepatocellular carcinoma (3,39,40). Endothelial dysfunction plays a critical role in the pathology of NAFLD and NASH (41). Indeed, liver sinusoidal endothelial cells (LSECs) can function to provoke inflammation and fibrosis in NAFLD (16). To delve further into the function of miR-494 during ER stress *in vivo*, we utilized a well characterized murine model of NASH (42). It has been shown that injection of TCN (2μg/g) increases acute ER stress in the liver with upregulation of CHOP and spliced XBP1 (42). We observed that TCN treatment induced endogenous miR-494 expression in mouse livers (Fig. 4A). To address the relationship between miR-494 and ER stress, we used hydrodynamic injection of plasmid DNA to transiently increase miR-494 levels in the liver. Compared to the GFP control plasmid, miR-494 plasmid injection significantly increased the levels of miR-494 (Fig. 4B). Importantly, when followed by TCN treatment 24h later, this increase in miR-494 levels was sufficient to decrease ER stress in the livers as assessed by *DDIT3* levels (Fig. 4C).

**Figure 4:**
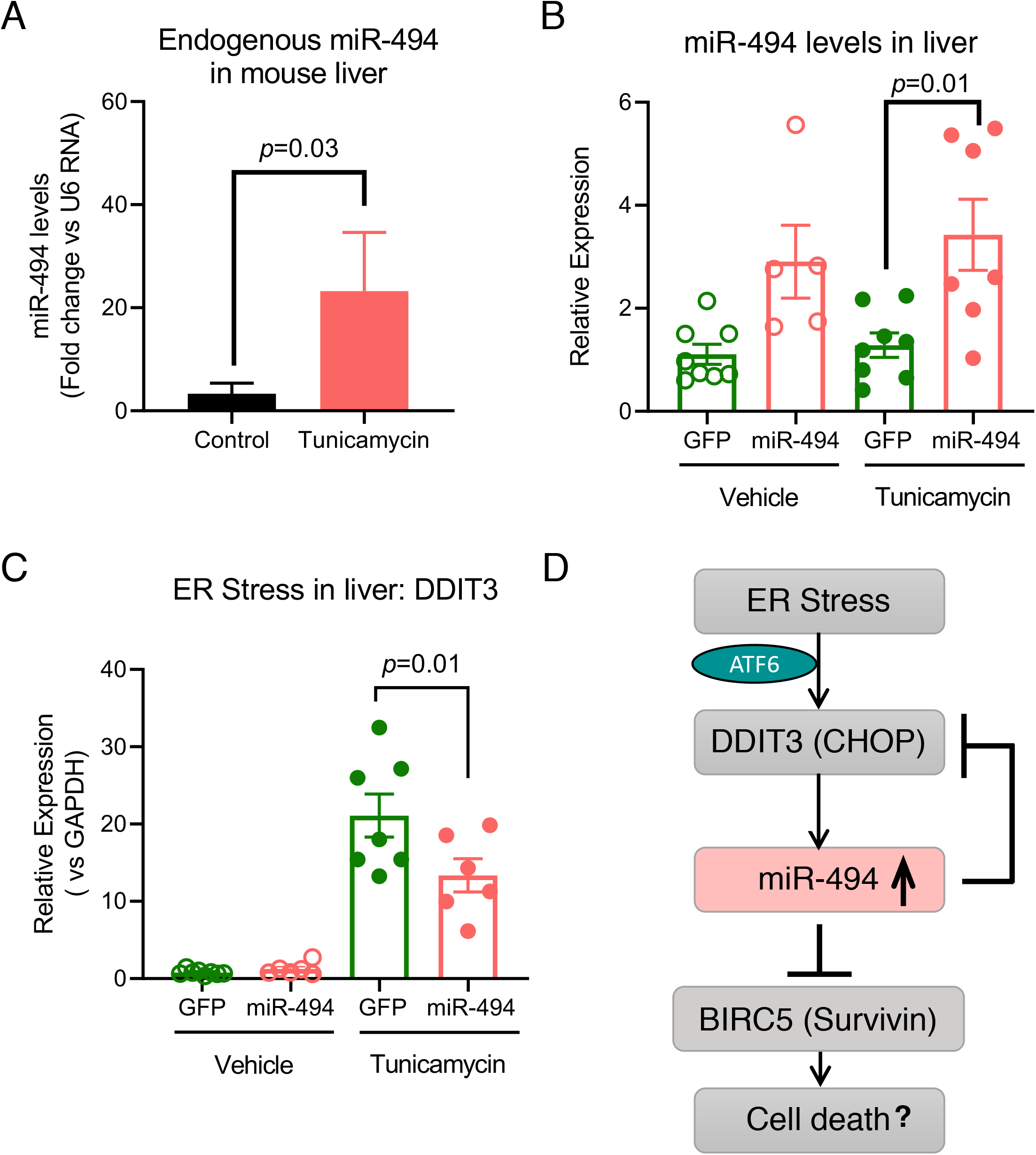
miR-494 regulates ER stress *in vivo*. miR-494 levels measured by qRT-PCR from whole liver harvested from male C57BL/6 mice injected *i.p.* with TCN (2μg/g, *n*=11) or vehicle control (*n*=10) for 6h. Bars show mean + SEM. P-values are from a Mann-Whitney U-test. B) miR-494 levels and C) ER stress levels measured by DDIT3 mRNA were assessed by qRT-PCR from whole livers harvested from male C57BL/6 mice (*n*=5-8 per group) after hydrodynamic injection of miR-494 or GFP control plasmids (20-30μg/animal, 48h) followed by TCN or vehicle treatment (24h). Livers of mice treated with Tunicamycin 24h after hydrodynamic injection of miR-494 or a control GFP plasmid. P=0.01 via one-way ANOVA with a Holm-Sidak’s post-hoc correction. Each dot represents an individual mouse. D) Schematic showing ER stress induction of miR-494 targets *BIRC5* and functions to dampen ER stress.

ER stress induction by TCN treatment has been shown to induce inflammation in the liver, which is a hallmark of NASH (3,7). To explore the mechanisms by which miR-494 may impact the activity of TCN *in vivo*, we utilized a Nanostring^®^ IO360 mouse profiling array to evaluate the immune response and inflammation in these animals. Compared to the plasmid control, miR-494 treated mice exhibited a significant decrease in several immune response and inflammatory genes in the liver (Supplementary Fig. 5A). Gene Ontology analysis identified cytokine response, immune cell migration and immune cell adhesion to vascular endothelium as the most significantly downregulated pathways in the miR-494 treatment group (Supplementary Fig. 5B).

## Discussion

Here we show that endoplasmic reticulum (ER) stress is a potent inducer of miR-494 in endothelial cells (ECs) *in vitro* and in murine liver *in vivo*. Our data indicates that the ER stress-associated increase in expression of miR-494 occurs in an ATF6 dependent manner. Upon induction, miR-494 decreases the level of survivin (*BIRC5*) —an anti-apoptotic protein— amongst other genes important in DNA replication and cell proliferation. We show that miR-494 gain-of-function diminishes the magnitude of the unfolded protein response (UPR) induced by tunicamycin (TCN) in both *in vitro* and *in vivo* models of acute ER stress. Functionally, mice pre-treated with a miR-494 overexpression plasmid exhibit a decrease in numerous inflammatory genes and cytokines in response to TCN in the liver. This miR-494 mediated decrease in ER stress appears to impact inflammation. Overall, this report shows that miR-494 downregulates critical targets in cell proliferation and survival which function to dampen the ER stress response by mechanisms which have yet to be revealed (Fig. 4D).

Many chronic human diseases have an established link within the ER stress response signaling cascades. Mechanistically, ER stress and the unfolded protein response (UPR) are triggered by the accumulation of misfolded proteins (2,43). The UPR is initiated by the dissociation of the molecular chaperone protein BiP from the three sensors, PERK, IRE1α and ATF6. Dissociation of BiP activates these proteins resulting either oligomerization or export from the ER. These activated sensors transduce signals to initiate pathways which work in a coordinated manner to halt protein translation, increase ER chaperone levels and clear misfolded proteins. Known chemical inducers of ER stress, TCN and thapsigargin (TG) generate a robust UPR in multiple models of disease including atherosclerosis and NASH (42,44). TCN is a toxic aminoglycoside antibiotic which triggers ER stress by preventing core oligosaccharide addition to nascent polypeptides, thereby blocking proper protein folding. Numerous studies have shown that TCN stimulates all three of the UPR signaling pathways causing an upsurge in transcription factor activity (*eg*. CHOP, XBP1 splicing), global inhibition of translation (eIF2α phosphorylation), and ATF6 translocation to the Golgi apparatus. We found that TCN treatment of ECs induced CHOP transcription and XBP1 splicing, which are indicative of an appropriate ER stress response. Further, TCN consistently generated a rapid and robust upregulation of miR-494 in multiple EC cell lines (Fig. 1 and Supplementary Fig. 2). TG, a non-competitive inhibitor of the Sarco/Endoplasmic Reticulum Calcium ATPase (SERCA) disrupts calcium homeostasis in the ER. In our experiments, TG treated ECs experienced similar signaling patterns, with an increase in both spliced *XBP1*, CHOP transcription and induction of miR-494 (Supplementary Fig 1). Interestingly, TCN but not TG induced a biphasic expression of mature miR-494 with a slight increase at 1h and a more significant (~30 fold) increase at 6h. Moreover, primary miR-494 transcript expression mirrored that of the mature transcript in TG treatment but not TCN (Fig. 1 & Supplementary Fig.1). It is unclear if these differences reflect differential regulation of miR processing pathways by these two inducers. Indeed, the canonical miR processing protein Dicer has been shown to localize and interact with proteins in the ER (45). Therefore, it is possible that ER stress pathways can differentially impact miR transcription and processing and it will be pertinent to investigate the impact of ER stress in the setting of EC-associated induction of miR-494.

Our data from gain-and loss-of-function experiments show that miR-494 decreases ER stress in ECs and in mouse liver (Fig. 2, 4). These results bolster other reports which also demonstrate miR-mediated responses to ER stress in different tissues (26,46–49). While miRs have a recognized role in liver disease (50), our studies are the first to show the ER stress reducing activity of miR-494 in the context of liver disease (NASH). miR-494 is known to be oncogenic and drives tumor progression and drug resistance in specific cancer types including colorectal cancer (51) and hepatocellular carcinoma (52,53). Furthermore, miR-494 has been shown to attenuate ischemia reperfusion injury in the liver by regulating the PI3K/Akt signaling pathway (54). We previously reported a unique role for miR-494 as a mediator of endothelial senescence by decreasing the MRN DNA repair protein complex. Other studies have shown that miR-494 is involved in vascular inflammation in atherosclerosis (55). Our data indicates a novel function of miR-494 in a complex pathological process.

Using a proteomics approach, we identified six putative miR-494 targets that are relevant during the ER stress response in ECs. Of these, survivin (*BIRC5*) has been previously shown to be a bona fide target of miR-494 in different disease models (37,38). Interestingly, the cross-talk between ER stress and survivin has been shown to downregulate inflammatory genes in a mouse model of chronic ER stress in the colon (56). Consistent with our observations of a miR-mediated reduction in ER stress combined with a decrease in survivin (*BIRC5*) and markers of inflammation, Gundamaraju *et al*., demonstrated that pharmacological inhibition of survivin is comparable to ER stress inhibition and attenuates inflammatory gene expression (56).

We used a Nanostring Mouse PanCancer™ profiling array to query the *in vivo* response to miR-494 pre-treatment prior to induction of ER stress. Our results reveal that a number of inflammatory genes are downregulated in the liver with miR-494 plasmid pre-treatment. Very few of the array-identified genes harbor miR-494 binding sites and therefore, the mechanism(s) by which miR-494 diminishes ER stress and subsequent inflammation is unclear. Further mechanistic complexity arises from recent reports which indicate miR-494 can be found in exosomes from patients with cirrhosis (57,58). Pertinent to this body of work, the relative contributions of miR-494 from hepatocytes in comparison to liver ECs requires further investigation. In this setting, it is conceivable that exosomal miR-494 from sinusoidal or other ECs in the liver can orchestrate inflammation and mediate paracrine effects during ER stress.

In summary, our work shows that miR-494 is induced during acute ER stress and functions to attenuate ER stress *in vitro* and *in vivo*. Our observations elucidate a new potential mechanistic role for miR-494 in the ER stress response pathway in ECs and likely in other cell types. These studies offer opportunities to inspire new hypotheses to understand the link between miR activity and the response to stressors which influence cell fate decisions in many human diseases.

---

**Supplementary Figure 1:**
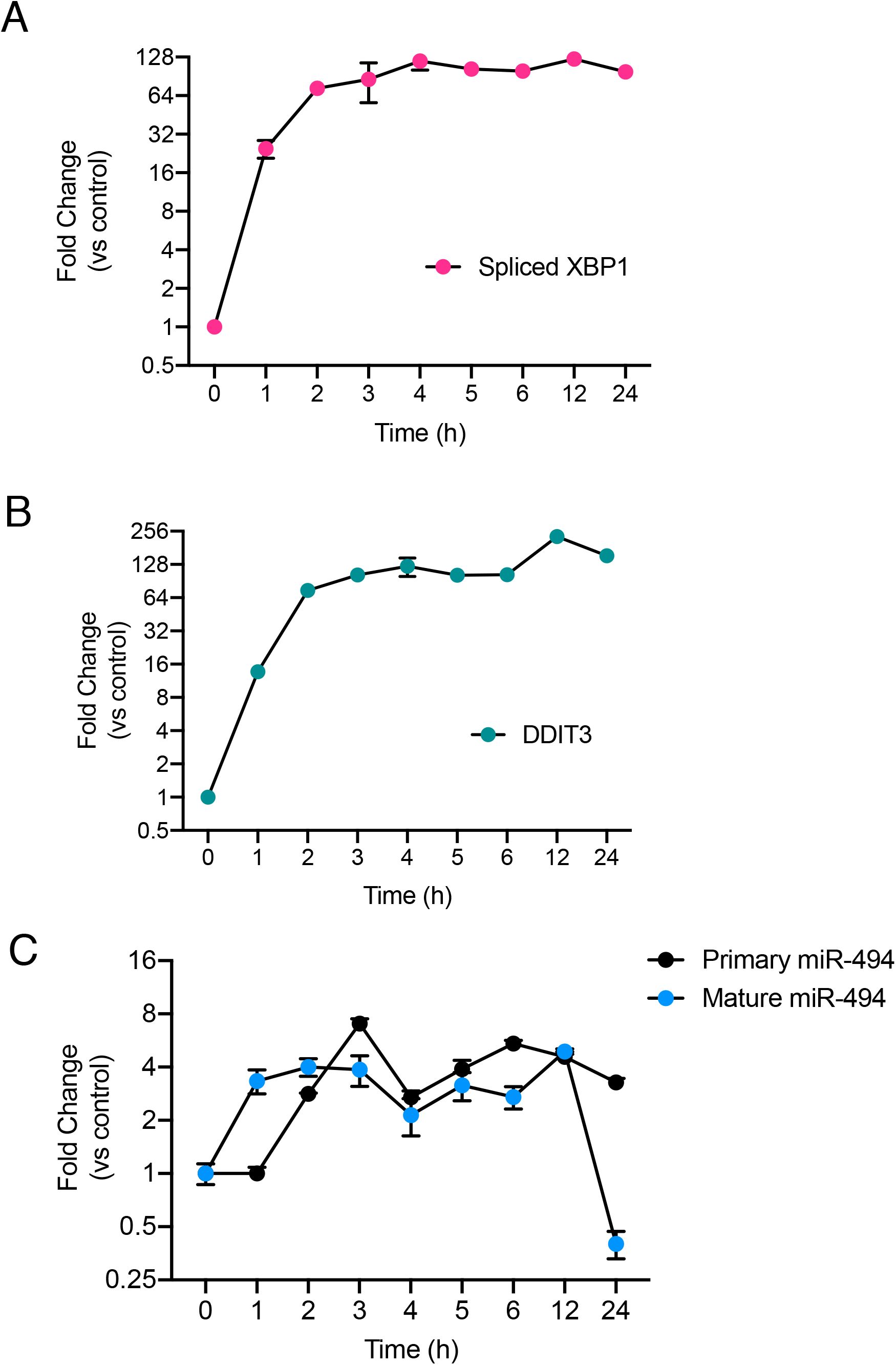
Thapsigargin induced ER stress drives miR-494 expression. Relative mRNA expression of A) *sXBP1* and B) *DDIT3* (CHOP) mRNAs and C) primary and mature miR-494 in HUVECs treated with 0.1μM Thapsigargin. Gene expression is normalized to GAPDH or U6 and mean fold change compared to vehicle control or time 0h is shown. Graphs are representative of 1 of 3 biological replicates. Values indicate mean ± standard deviation.

**Supplementary Figure 2:**
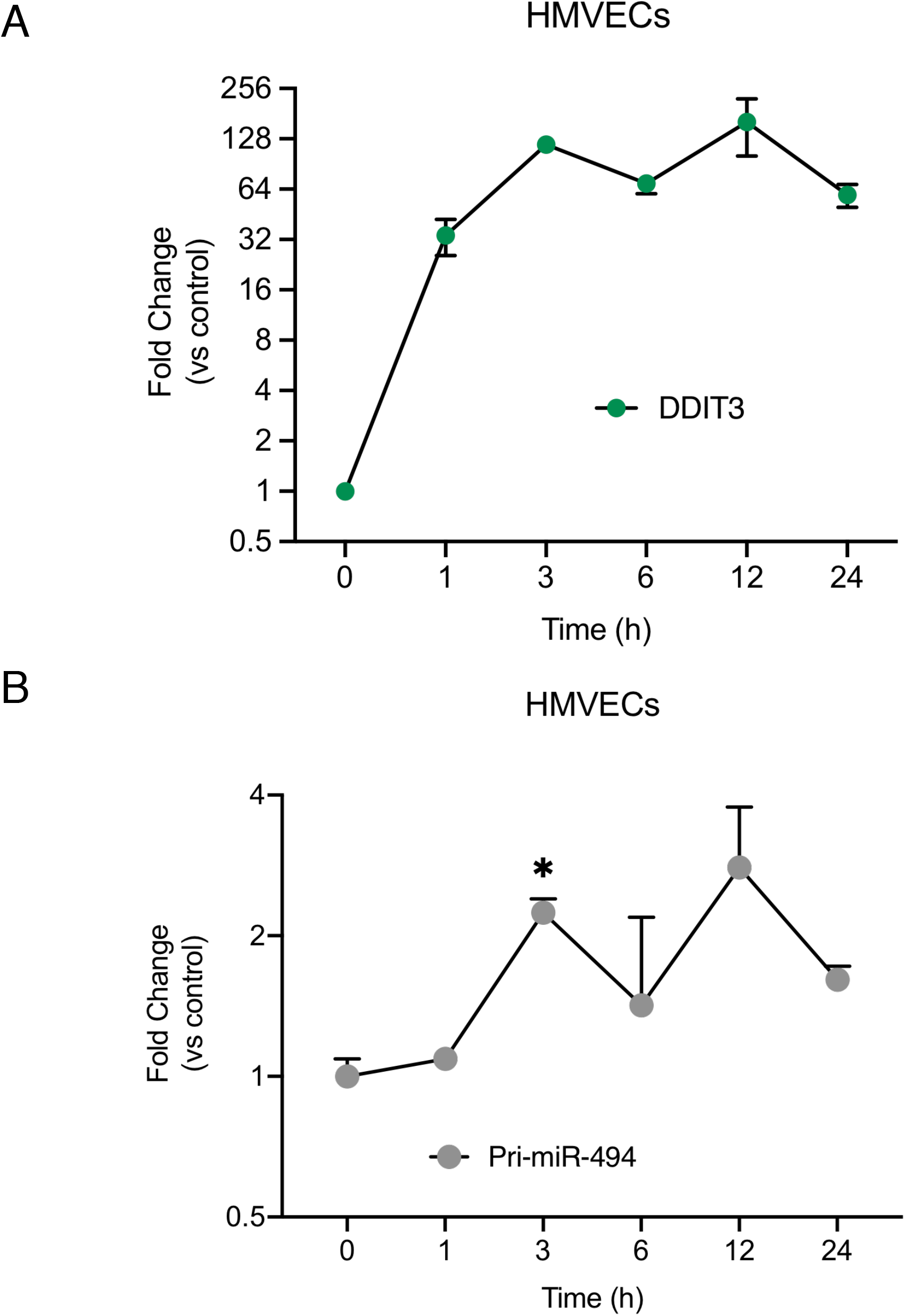
ER stress induces expression of miR-494 in human microvascular ECs. A. Relative mRNA expression of *DDIT3* (CHOP) in HMVECs treated with 5μg/mL Tunicamycin. (A) Relative expression of primary miR-494 treated with Tunicamycin. Gene expression is normalized to GAPDH and mean fold change compared to vehicle control or time 0h is shown. Graphs are representative of 1 biological replicate from 3 independent replicates where values indicate mean ± standard deviation. * P<0.05 using two-way ANOVA with Fisher’s LSD test.

**Supplementary Figure 3:**
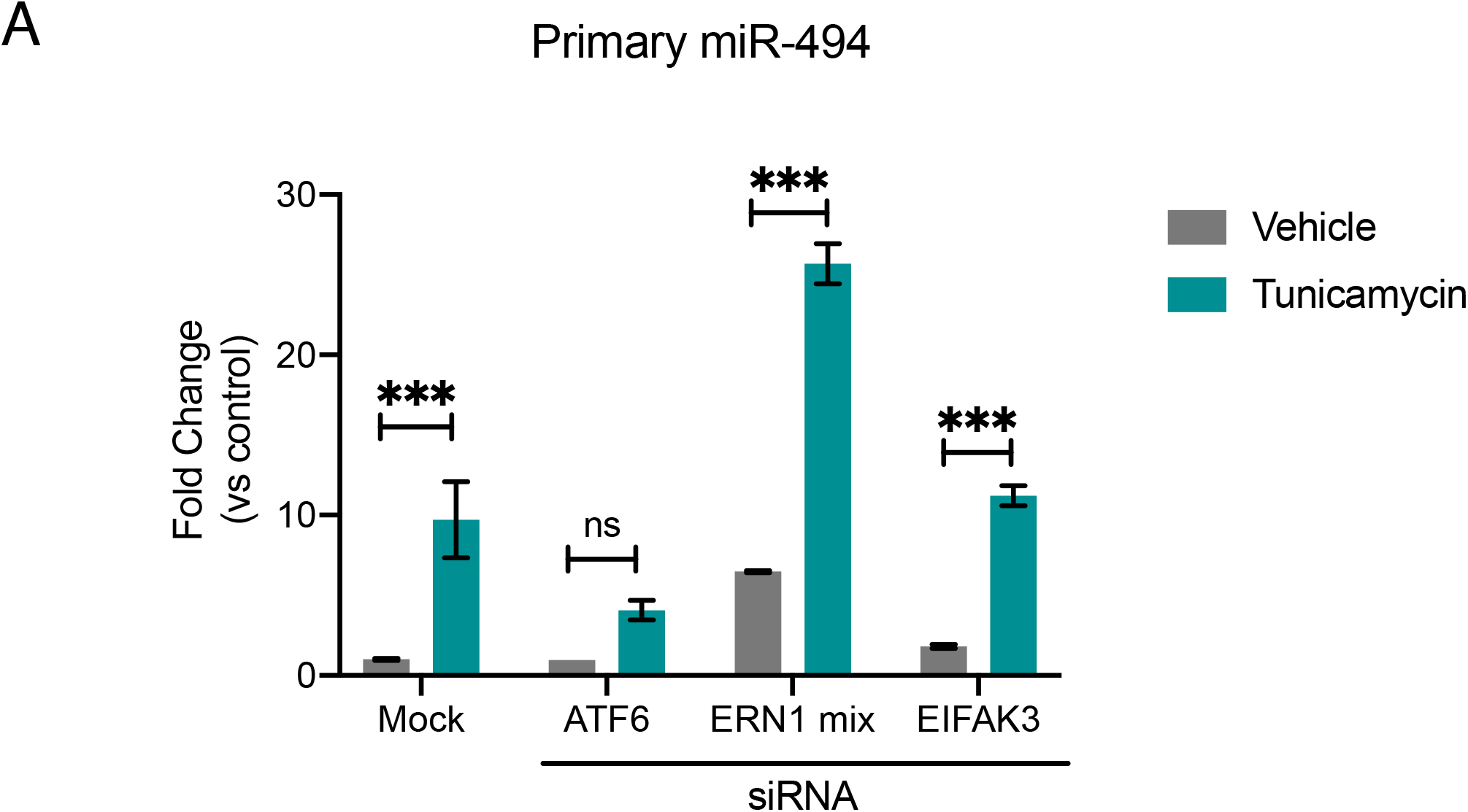
ER stress via ATF6 induces expression of miR-494. HUVECs were transfected with indicated siRNAs. After 48h, cells were treated with vehicle or Tunicamycin (10μg/mL). Relative expression of primary miR-494 normalized to GAPDH is shown. Mean fold change compared to vehicle control is shown. Graphs are representative of 1 biological replicate from 2 independent replicates where values indicate mean ± standard deviation. * *p*<0.001 using two-tailed Student’s T-test.

**Supplementary Figure 4:**
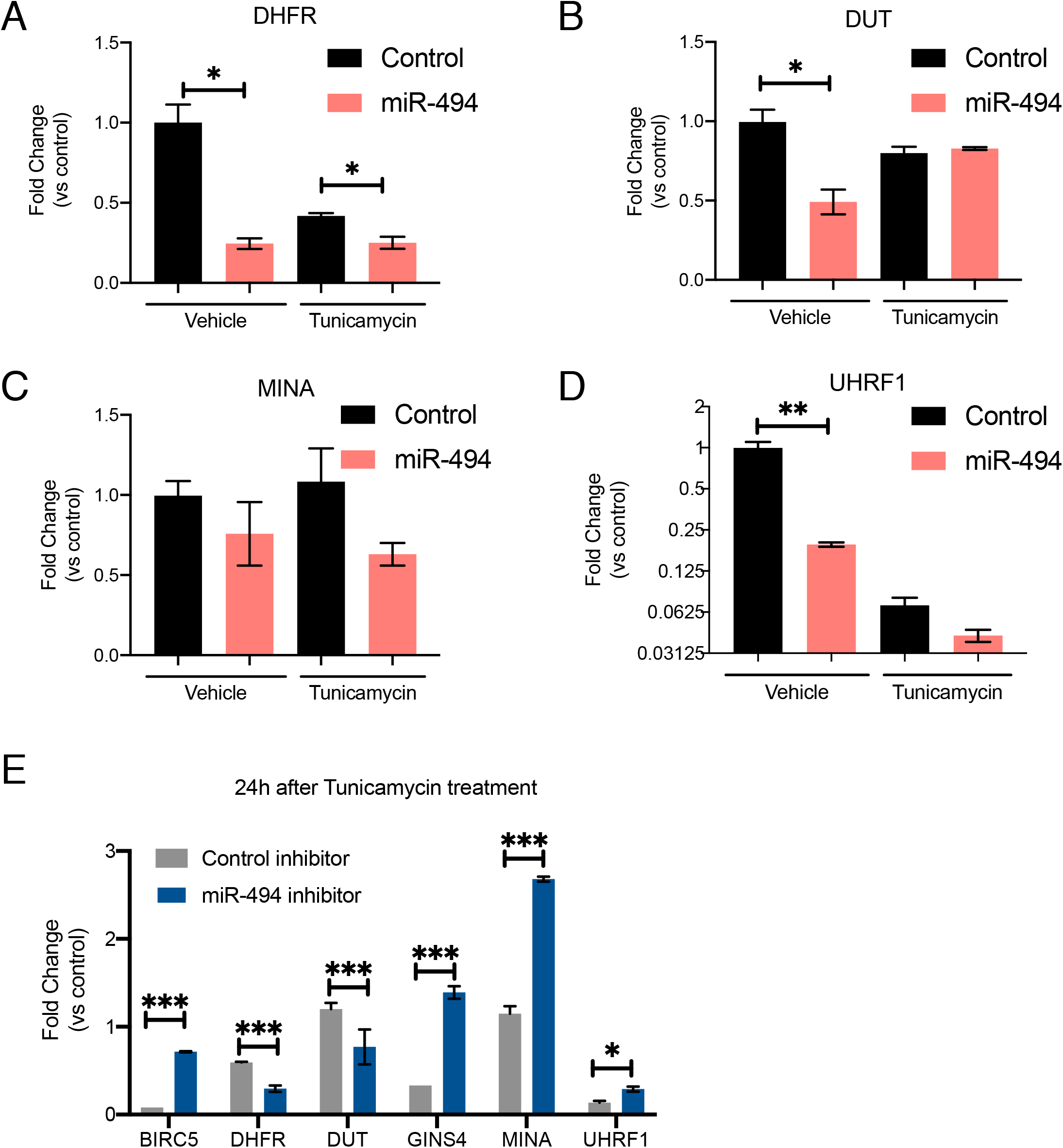
miR-494 regulates several target genes in cell survival and DNA replication. A-D) Fold-change (compared to respective controls) of mRNA levels as assessed by qRT-PCR for the 4 targets that are downregulated in both Tunicamycin and miR-494 groups in Fig 3B. HUVECs were transfected and treated as described in 2A. Gene expression measured iat 48h post treatment. E) HUVECs were transfected with either a control inhibitor or miR-494 inhibitor as shown in 2B. mRNA levels of indicated target genes were evaluated using qRT-PCR. Bars show mean + S.D. of one representative experiment out of three independent biological replicates. * P<0.05, ** P< 0.01, ***P<0.001 by two-tailed Student’s T-tests comparing respective control groups.

**Supplementary Figure 5:**
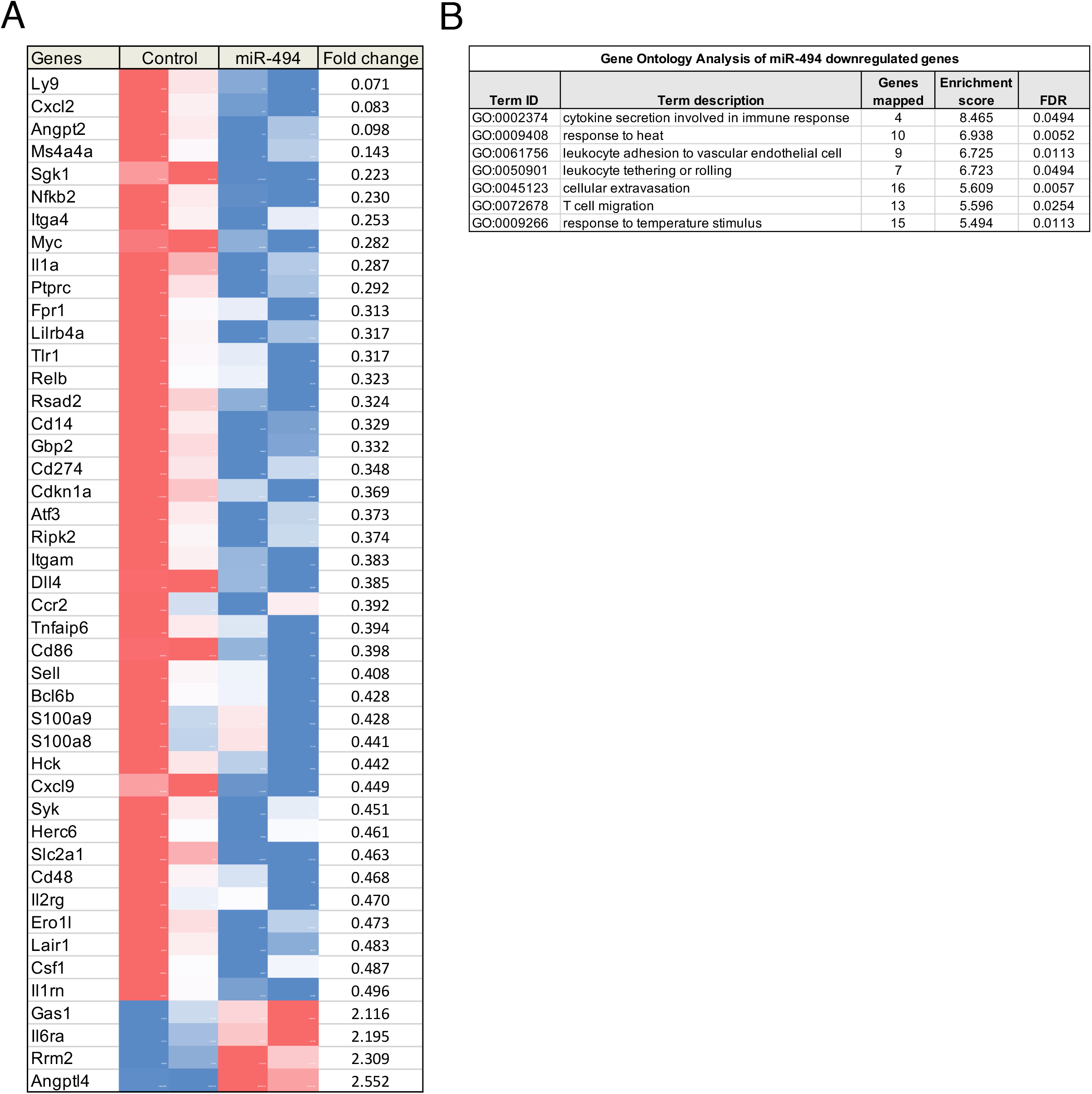
miR-494 treatment *in vivo* impacts inflammatory gene expression. A) Heatmap depicting normalized gene expression in mice treated with miR-494 plasmid (n=2) or a control plasmid (n=2) via hydrodynamic injection followed by Tunicamycin treatment from experiment shown in Fig 4B-C. Gene expression was assessed using a Nanostring IO360 panel containing 770 tumor microenvironment, immune response and inflammation genes. Heatmap depicts genes at least 2 fold downregulated or upregulated compared to control plasmid. B) Gene Ontology analysis based on the whole 770 gene panel using STRING database. All the GO enriched genes derived from the miR-494 downregulated gene cluster.

## References

1. Ellgaard, L., and Helenius, A. (2003) Quality control in the endoplasmic reticulum. Nature Reviews Molecular Cell Biology 4, 181–191

2. Ron, D., and Walter, P. (2007) Signal integration in the endoplasmic reticulum unfolded protein response. Nature Reviews Molecular Cell Biology 8, 519–529

3. Malhi, H., and Kaufman, R. J. (2011) Endoplasmic reticulum stress in liver disease. Journal of Hepatology 54, 795–809

4. Zhang, K., and Kaufman, R. J. (2008) From endoplasmic-reticulum stress to the inflammatory response. Nature 454, 455–462

5. Brodsky, S. V., and Goligorsky, M. S. (2012) Endothelium Under Stress: Local and Systemic Messages. Seminars in Nephrology 32, 192–198

6. Lenna, S., Han, R., and Trojanowska, M. (2014) Endoplasmic reticulum stress and endothelial dysfunction. IUBMB Life 66, 530–537

7. Gargalovic, P. S., Gharavi, N. M., Clark, M. J., Pagnon, J., Yang, W.-P., He, A., Truong, A., Baruch-Oren, T., Berliner, J. A., Kirchgessner, T. G., and Lusis, A. J. (2006) The Unfolded Protein Response Is an Important Regulator of Inflammatory Genes in Endothelial Cells. Arteriosclerosis, Thrombosis, and Vascular Biology 26, 2490–2496

8. Tabas, I. (2010) The role of endoplasmic reticulum stress in the progression of atherosclerosis. Circ Res 107, 839–850

9. Civelek, M., Manduchi, E., Riley, R. J., Stoeckert, C. J., Jr., and Davies, P. F. (2009) Chronic endoplasmic reticulum stress activates unfolded protein response in arterial endothelium in regions of susceptibility to atherosclerosis. Circ Res 105, 453–461

10. Zeng, L., Zampetaki, A., Margariti, A., Pepe, A. E., Alam, S., Martin, D., Xiao, Q., Wang, W., Jin, Z. G., Cockerill, G., Mori, K., Li, Y. S., Hu, Y., Chien, S., and Xu, Q. (2009) Sustained activation of XBP1 splicing leads to endothelial apoptosis and atherosclerosis development in response to disturbed flow. Proc Natl Acad Sci U S A 106, 8326–8331

11. Kassan, M., Galán, M., Partyka, M., Saifudeen, Z., Henrion, D., Trebak, M., and Matrougui, K. (2012) Endoplasmic Reticulum Stress Is Involved in Cardiac Damage and Vascular Endothelial Dysfunction in Hypertensive Mice. Arteriosclerosis, Thrombosis, and Vascular Biology 32, 1652–1661

12. R.J., C. S. S. a. K. (2014) Endoplasmic Reticulum Stress and Oxidative Stress in Cell Fate Decision and Human Disease. Antioxidants & Redox Signaling 21, 396–413

13. Maamoun, H., Abdelsalam, S. S., Zeidan, A., Korashy, H. M., and Agouni, A. (2019) Endoplasmic Reticulum Stress: A Critical Molecular Driver of Endothelial Dysfunction and Cardiovascular Disturbances Associated with Diabetes. Int J Mol Sci 20

14. Maamoun, H., Benameur, T., Pintus, G., Munusamy, S., and Agouni, A. (2019) Crosstalk Between Oxidative Stress and Endoplasmic Reticulum (ER) Stress in Endothelial Dysfunction and Aberrant Angiogenesis Associated With Diabetes: A Focus on the Protective Roles of Heme Oxygenase (HO)-1. Frontiers in Physiology 10

15. Zhou, Q. G., Fu, X. J., Xu, G. Y., Cao, W., Liu, H. F., Nie, J., Liang, M., and Hou, F. F. (2012) Vascular insulin resistance related to endoplasmic reticulum stress in aortas from a rat model of chronic kidney disease. American Journal of Physiology-Heart and Circulatory Physiology 303, H1154–H1165

16. Hammoutene, A., and Rautou, P.-E. (2019) Role of liver sinusoidal endothelial cells in non-alcoholic fatty liver disease. Journal of Hepatology 70, 1278–1291

17. Cao, S. S. a. K., R.J. (2014) Endoplasmic Reticulum Stress and Oxidative Stress in Cell Fate Decision and Human Disease. Antioxidants & Redox Signaling 21, 396–413

18. Leung, A. K. L., and Sharp, P. A. (2010) MicroRNA Functions in Stress Responses. Molecular Cell 40, 205–215

19. Olejniczak, M., Kotowska-Zimmer, A., and Krzyzosiak, W. (2018) Stress-induced changes in miRNA biogenesis and functioning. Cell Mol Life Sci 75, 177–191

20. Shi, L., Fisslthaler, B., Zippel, N., Frömel, T., Hu, J., Elgheznawy, A., Heide, H., Popp, R., and Fleming, I. (2013) MicroRNA-223 Antagonizes Angiogenesis by Targeting β1 Integrin and Preventing Growth Factor Signaling in Endothelial Cells. Circulation Research 113, 1320–1330

21. Wang, S., Aurora, A. B., Johnson, B. A., Qi, X., McAnally, J., Hill, J. A., Richardson, J. A., Bassel-Duby, R., and Olson, E. N. (2008) The Endothelial-Specific MicroRNA miR-126 Governs Vascular Integrity and Angiogenesis. Developmental Cell 15, 261–271

22. Ghosh, G., Subramanian, I. V., Adhikari, N., Zhang, X., Joshi, H. P., Basi, D., Chandrashekhar, Y. S., Hall, J. L., Roy, S., Zeng, Y., and Ramakrishnan, S. (2010) Hypoxia-induced microRNA-424 expression in human endothelial cells regulates HIF-α isoforms and promotes angiogenesis. The Journal of Clinical Investigation 120, 4141–4154

23. Voellenkle, C., van Rooij, J., Guffanti, A., Brini, E., Fasanaro, P., Isaia, E., Croft, L., David, M., Capogrossi, M. C., Moles, A., Felsani, A., and Martelli, F. (2012) Deep-sequencing of endothelial cells exposed to hypoxia reveals the complexity of known and novel microRNAs. RNA 18, 472–484

24. Espinosa-Diez, C., Wilson, R., Chatterjee, N., Hudson, C., Ruhl, R., Hipfinger, C., Helms, E., Khan, O. F., Anderson, D. G., and Anand, S. (2018) MicroRNA regulation of the MRN complex impacts DNA damage, cellular senescence, and angiogenic signaling. Cell Death Dis 9, 632

25. Chen, Z., Wen, L., Martin, M., Hsu, C.-Y., Fang, L., Lin, F.-M., Lin, T.-Y., Geary, M. J., Geary, G. G., Zhao, Y., Johnson, D. A., Chen, J.-W., Lin, S.-J., Chien, S., Huang, H.-D., Miller, Y. I., Huang, P.-H., and Shyy, J. Y.-J. (2015) Oxidative Stress Activates Endothelial Innate Immunity via Sterol Regulatory Element Binding Protein 2 (SREBP2) Transactivation of MicroRNA-92a. Circulation 131, 805–814

26. Maurel, M., and Chevet, E. (2013) Endoplasmic reticulum stress signaling: the microRNA connection. American Journal of Physiology-Cell Physiology 304, C1117–C1126

27. Upton, J. P., Wang, L., Han, D., Wang, E. S., Huskey, N. E., Lim, L., Truitt, M., McManus, M. T., Ruggero, D., Goga, A., Papa, F. R., and Oakes, S. A. (2012) IRE1alpha cleaves select microRNAs during ER stress to derepress translation of proapoptotic Caspase-2. Science 338, 818–822

28. Hiramatsu, N., Chiang, K., Aivati, C., Rodvold, J. J., Lee, J. M., Han, J., Chea, L., Zanetti, M., Koo, E. H., and Lin, J. H. (2020) PERK-mediated induction of microRNA-483 disrupts cellular ATP homeostasis during the unfolded protein response. J Biol Chem 295, 237–249

29. Sohn, E. J. (2018) (MicroRNA 200c-3p regulates autophagy via upregulation of endoplasmic reticulum stress in PC-3 cells). Cancer Cell Int 18, 2

30. Kassan, M., Vikram, A., Li, Q., Kim, Y.-R., Kumar, S., Gabani, M., Liu, J., Jacobs, J. S., and Irani, K. (2017) MicroRNA-204 promotes vascular endoplasmic reticulum stress and endothelial dysfunction by targeting Sirtuin1. Scientific Reports 7, 9308

31. Wilson, R., Espinosa-Diez, C., Kanner, N., Chatterjee, N., Ruhl, R., Hipfinger, C., Advani, S. J., Li, J., Khan, O. F., Franovic, A., Weis, S. M., Kumar, S., Coussens, L. M., Anderson, D. G., Chen, C. C., Cheresh, D. A., and Anand, S. (2016) MicroRNA regulation of endothelial TREX1 reprograms the tumour microenvironment. Nat Commun 7, 13597

32. Rana, S., Espinosa-Diez, C., Ruhl, R., Chatterjee, N., Hudson, C., Fraile-Bethencourt, E., Agarwal, A., Khou, S., Thomas, C. R., and Anand, S. (2020) Differential regulation of microRNA-15a by radiation affects angiogenesis and tumor growth via modulation of acid sphingomyelinase. Scientific Reports 10, 5581

33. Sakoda, Y., Anand, S., Zhao, Y., Park, J.-J., Liu, Y., Kuramasu, A., van Rooijen, N., Chen, L., Strome, S. E., Hancock, W. W., Chen, L., and Tamada, K. (2011) Herpesvirus entry mediator regulates hypoxia-inducible factor–1α and erythropoiesis in mice. The Journal of Clinical Investigation 121, 4810–4819

34. Bonamassa, B., Hai, L., and Liu, D. (2011) Hydrodynamic gene delivery and its applications in pharmaceutical research. Pharm Res 28, 694–701

35. Plubell, D. L., Wilmarth, P. A., Zhao, Y., Fenton, A. M., Minnier, J., Reddy, A. P., Klimek, J., Yang, X., David, L. L., and Pamir, N. (2017) Extended Multiplexing of Tandem Mass Tags (TMT) Labeling Reveals Age and High Fat Diet Specific Proteome Changes in Mouse Epididymal Adipose Tissue. Mol Cell Proteomics 16, 873–890

36. Xu, S., Li, D., Li, T., Qiao, L., Li, K., Guo, L., and Liu, Y. (2018) miR-494 Sensitizes Gastric Cancer Cells to TRAIL Treatment Through Downregulation of Survivin. Cellular physiology and biochemistry : international journal of experimental cellular physiology, biochemistry, and pharmacology 51, 2212–2223

37. Yun, S., Kim, W. K., Kwon, Y., Jang, M., Bauer, S., and Kim, H. (2018) Survivin is a novel transcription regulator of KIT and is downregulated by miRNA-494 in gastrointestinal stromal tumors. International journal of cancer 142, 2080–2093

38. Zhu, J., Sun, C., Wang, L., Xu, M., Zang, Y., Zhou, Y., Liu, X., Tao, W., Xue, B., Shan, Y., and Yang, D. (2015) Targeting survivin using a combination of miR-494 and survivin shRNA has synergistic effects on the suppression of prostate cancer growth. Molecular medicine reports 13

39. Maiers, J. L., and Malhi, H. (2019) Endoplasmic Reticulum Stress in Metabolic Liver Diseases and Hepatic Fibrosis. Seminars in liver disease 39, 235–248

40. Lebeaupin, C., Vallée, D., Hazari, Y., Hetz, C., Chevet, E., and Bailly-Maitre, B. (2018) Endoplasmic reticulum stress signalling and the pathogenesis of non-alcoholic fatty liver disease. J Hepatol 69, 927–947

41. Pasarín, M., Abraldes, J. G., Liguori, E., Kok, B., and La Mura, V. (2017) Intrahepatic vascular changes in non-alcoholic fatty liver disease: Potential role of insulin-resistance and endothelial dysfunction. World J Gastroenterol 23, 6777–6787

42. Lee, J. S., Zheng, Z., Mendez, R., Ha, S. W., Xie, Y., and Zhang, K. (2012) Pharmacologic ER stress induces non-alcoholic steatohepatitis in an animal model. Toxicology letters 211, 29–38

43. Wang, M., and Kaufman, R. J. (2016) Protein misfolding in the endoplasmic reticulum as a conduit to human disease. Nature 529, 326–335

44. Witte, I., and Horke, S. (2011) Assessment of endoplasmic reticulum stress and the unfolded protein response in endothelial cells. Methods in enzymology 489, 127–146

45. Pépin, G., Perron, M. P., and Provost, P. (2012) Regulation of human Dicer by the resident ER membrane protein CLIMP-63. Nucleic acids research 40, 11603–11617

46. Yang, F., Zhang, L., Wang, F., Wang, Y., Huo, X. S., Yin, Y. X., Wang, Y. Q., Zhang, L., and Sun, S. H. (2011) Modulation of the unfolded protein response is the core of microRNA-122-involved sensitivity to chemotherapy in hepatocellular carcinoma. Neoplasia (New York, N.Y.) 13, 590–600

47. Byrd, A. E., Aragon, I. V., and Brewer, J. W. (2012) MicroRNA-30c-2* limits expression of proadaptive factor XBP1 in the unfolded protein response. Journal of Cell Biology 196, 689–698

48. Belmont, P. J., Chen, W. J., Thuerauf, D. J., and Glembotski, C. C. (2012) Regulation of microRNA expression in the heart by the ATF6 branch of the ER stress response. Journal of Molecular and Cellular Cardiology 52, 1176–1182

49. Behrman, S., Acosta-Alvear, D., and Walter, P. (2011) A CHOP-regulated microRNA controls rhodopsin expression. The Journal of cell biology 192, 919–927

50. Szabo, G., and Bala, S. (2013) MicroRNAs in liver disease. Nature Reviews Gastroenterology & Hepatology 10, 542–552

51. Zhang, Y., Guo, L., Li, Y., Feng, G.-H., Teng, F., Li, W., and Zhou, Q. (2018) MicroRNA-494 promotes cancer progression and targets adenomatous polyposis coli in colorectal cancer. Molecular Cancer 17, 1

52. Zhang, J., Zhu, Y., Hu, L., Yan, F., and Chen, J. (2019) miR-494 induces EndMT and promotes the development of HCC (Hepatocellular Carcinoma) by targeting SIRT3/TGF-β/SMAD signaling pathway. Scientific Reports 9, 7213

53. Pollutri, D., Patrizi, C., Marinelli, S., Giovannini, C., Trombetta, E., Giannone, F. A., Baldassarre, M., Quarta, S., Vandewynckel, Y. P., Vandierendonck, A., Van Vlierberghe, H., Porretti, L., Negrini, M., Bolondi, L., Gramantieri, L., and Fornari, F. (2018) The epigenetically regulated miR-494 associates with stem-cell phenotype and induces sorafenib resistance in hepatocellular carcinoma. Cell Death & Disease 9, 4

54. Su, S., Luo, D., Liu, X., Liu, J., Peng, F., Fang, C., and Li, B. (2017) miR-494 up-regulates the PI3K/Akt pathway via targetting PTEN and attenuates hepatic ischemia/reperfusion injury in a rat model. Bioscience Reports 37

55. van Ingen, E., Foks, A. C., Kröner, M. J., Kuiper, J., Quax, P. H. A., Bot, I., and Nossent, A. Y. (2019) Antisense Oligonucleotide Inhibition of MicroRNA-494 Halts Atherosclerotic Plaque Progression and Promotes Plaque Stabilization. Molecular Therapy - Nucleic Acids 18, 638–649

56. Gundamaraju, R., Vemuri, R., Chong, W. C., Myers, S., Norouzi, S., Shastri, M. D., and Eri, R. (2018) Interplay between Endoplasmic Reticular Stress and Survivin in Colonic Epithelial Cells. Cells 7, 171

57. Li, J., Chen, J., Wang, S., Li, P., Zheng, C., Zhou, X., Tao, Y., Chen, X., Sun, L., Wang, A., Cao, K., Tang, S., and Zhou, J. (2019) Blockage of transferred exosome-shuttled miR-494 inhibits melanoma growth and metastasis. Journal of Cellular Physiology 234, 15763–15774

58. Fornari, F., Ferracin, M., Trerè, D., Milazzo, M., Marinelli, S., Galassi, M., Venerandi, L., Pollutri, D., Patrizi, C., Borghi, A., Foschi, F. G., Stefanini, G. F., Negrini, M., Bolondi, L., and Gramantieri, L. (2015) Circulating microRNAs, miR-939, miR-595, miR-519d and miR-494, Identify Cirrhotic Patients with HCC. PLOS ONE 10, e0141448

